# Genome-wide fitness assessment during diurnal growth reveals an expanded role of the cyanobacterial circadian clock protein KaiA

**DOI:** 10.1101/283812

**Authors:** David G. Welkie, Benjamin E. Rubin, Yong-Gang Chang, Spencer Diamond, Scott A. Rifkin, Andy LiWang, Susan S. Golden

## Abstract

The recurrent pattern of light and darkness generated by Earth’s axial rotation has profoundly influenced the evolution of organisms, selecting for both biological mechanisms that respond acutely to environmental changes and circadian clocks that program physiology in anticipation of daily variations. The necessity to integrate environmental responsiveness and circadian programming is exemplified in photosynthetic organisms such as cyanobacteria, which depend on light-driven photochemical processes. The cyanobacterium *Synechococcus elongatus* PCC 7942 is an excellent model system for dissecting these entwined mechanisms. Its core circadian oscillator, consisting of three proteins KaiA, KaiB, and KaiC, transmits time-of-day signals to clock-output proteins, which reciprocally regulate global transcription. Research performed under constant light facilitates analysis of intrinsic cycles separately from direct environmental responses, but does not provide insight into how these regulatory systems are integrated during light-dark cycles. Thus, we sought to identify genes that are specifically necessary in a day-night environment. We screened a dense bar-coded transposon library in both continuous light and daily cycling conditions and compared the fitness consequences of loss of each nonessential gene in the genome. Although the clock itself is not essential for viability in light-dark cycles, the most detrimental mutations revealed by the screen were those that disrupt KaiA. The screen broadened our understanding of light-dark survival in photosynthetic organisms, identified unforeseen clock-protein interaction dynamics, and reinforced the role of the clock as a negative regulator of a night-time metabolic program that is essential for *S. elongatus* to survive in the dark.

**Significance:** Understanding how photosynthetic bacteria respond to and anticipate natural light–dark cycles is necessary for predictive modeling, bioengineering, and elucidating metabolic strategies for diurnal growth. Here, we identify the genetic components that are important specifically under light-dark cycling conditions and determine how a properly functioning circadian clock prepares metabolism for darkness, a starvation period for photoautotrophs. This study establishes that the core circadian clock protein KaiA is necessary to enable rhythmic de-repression of a nighttime circadian program.

**Specific Contributions:** DGW BR SD and SSG conceived and designed the project. DGW BR and YGC performed the experiments and analyzed the data. SAR wrote the R scripts for RB-TnSeq conditional fitness analysis. YGC and AL interpreted fluorescence anisotropy data. DGW BR SD SSG YGC and AL wrote the manuscript.

## Introduction

Photosynthetic organisms experience a dramatic change in physiology and metabolism each day when the sun sets and the daytime processes related to photosynthesis become inoperative. Like many diverse organisms throughout nature, prokaryotic cyanobacteria have evolved circadian clocks that aid in the temporal orchestration of activities that are day-or night-appropriate. The circadian clock’s contribution towards fitness in changing environments was first demonstrated in 1998, when it was shown that strains of the cyanobacterium *Synechococcus elongatus* PCC 7942 with intrinsic circadian periods that match that of external light-dark cycles (LDC) outcompete strains that have different periods (1, 2). More recent experiments showed the value of preparing for night: when cells synchronized to different phases of the circadian cycle are mixed and then exposed to a pulse of darkness, those whose internal clock corresponds to a dawn or day-time phase have a higher occurrence of arrested growth upon re-illumination than those synchronized to anticipate darkness at the time of the pulse (3). These data support the hypothesis that the clock acts to prepare the cell for conditions that are predictable and recurrent, such as night following day.

A fitness advantage of appropriate circadian timing signals is not exclusive to cyanobacteria, as is observed in the model plant *Arabidopsis* (4, 5) and in humans, where circadian disruption can result in myriad adverse health effects including cardiovascular disease, cancer, and sleep disorders (6–8). Although the fundamental properties of circadian rhythms are shared across the domains of life, the molecular networks that generate them vary (9). In *S. elongatus* a three-protein oscillator comprising the proteins KaiA, KaiB, and KaiC sets up a circadian cycle of KaiC phosphorylation. This oscillator even functions *in vitro* when mixed at appropriate ratios with ATP (10), which has provided a powerful tool for exploring the mechanism that generates circadian rhythms. Genetic, biochemical, metabolic, and structural inquiries have revealed that the *S. elongatus* oscillator actively turns off a night-time metabolic program prior to dawn by triggering the dephosphorylation of the master transcription factor Regulator of Phycobilisome Association A (RpaA). It accomplishes this task by recruiting Circadian Input Kinase A (CikA) to a ring of KaiB that forms on KaiC during the night-time portion of the cycle, thus activating CikA phosphatase activity against RpaA (11–13). Among the targets of the RpaA regulon are the genes that facilitate catabolism of glycogen and generation of NADPH via the oxidative pentose phosphate pathway (14). These enzymatic reactions are essential for the cyanobacterium to survive the night when photosynthetic generation of reductant is disabled (15, 16).

While the molecular mechanisms of the circadian clock are well established in constant light for *S. elongatus* (17), its role in response to LDC, and the mechanisms by which the cell integrates circadian and environmental data, are still not understood. There have been few publications on the topic, and this study is the first to report on a genome-wide screen for LDC-sensitive mutants. Studies that report conditional LDC effects have been limited to a small number of genes. For instance, mutants defective for the circadian clock output pathway genes, *rpaA* and *sasA* (15, 16, 18, 19), and mutants unable to mobilize carbon through the oxidative pentose phosphate pathway (OPPP) and glycogen breakdown (16–20, 21), or unable to synthesize the alarmone nucleotide ppGpp (22, 23), have conditional defects specific to growth in LDC. To identify the genetic contributors to fitness under LDC and acquire a more comprehensive picture of the regulatory network, we used an unbiased population-based screen that tracks the fitness contributions of individual *S. elongatus* genes to reproduction in diel cycles. The method, random bar code transposon-site sequencing (RB-TnSeq), quantitatively compares the changes in abundances of individuals in a pooled mutant population over time. This assay allowed the calculation of a fitness contribution to LDC survival for each nonessential gene in the genome by leveraging multiple inactivating mutations spread across a genomic locus. This library was previously used to identify the essential gene set of *S. elongatus* under standard laboratory continuous light conditions (CLC) (24) and to generate an improved the metabolic model for generating phenotype predictions (25).

The results described here provide a comprehensive picture of genes that contribute to fitness under diurnal growth conditions. In addition to identifying the genetic components previously known to be critical for growth in LDC we found ~100 new genes whose mutation confers weak to strong fitness disadvantages. Of these genes, we focused on the top-scoring elements for LDC-specific defects which, unexpectedly, lie within the *kaiA* gene of the core circadian oscillator. As no previously known roles for KaiA predicted this outcome, we characterized the mechanism that underlies the LDC sensitivity of a kaiA-null mutant, and revealed interactions within the oscillator complex that regulate the activation of CikA, an RpaA phosphatase.

## Results

### Growth of library mutants to assess genetic fitness contributions under light-dark cycles

The RB-TnSeq mutant library includes a unique identifier sequence, or bar code, at each insertion site to track each transposon-insertion mutant. Next-generation sequencing is used to quantify these bar codes within the total DNA of the population, such that the survival and relative fitness of the approximately 150,000 mutants in the library can be tracked under control and experimental conditions (26). In this way, the *S. elongatus* RB-TnSeq library can be used for pooled, quantitative, whole-genome mutant screens.

First, the library was thawed, placed into 250-mL flasks, and sampled during incubation in CLC under only the selective pressure for the transposon antibiotic-resistance marker. Genomic DNA was extracted, and the bar codes of the mutant population were sequenced to establish a baseline abundance (Fig. 1). The culture was then divided into photobioreactors set to maintain constant optical density under either CLC, or LDC conditions that mimic day and night. After a treatment period allowing for approximately 6 generations, the library was again sampled and bar codes were quantified in a similar fashion. By comparing the frequencies of the bar codes after growth in LDC to those in CLC (<u>Dataset S1</u>), we estimated a quantitative measure of the fitness contribution for each gene specifically under LDC (**Dataset S2**).

**Fig 1.**
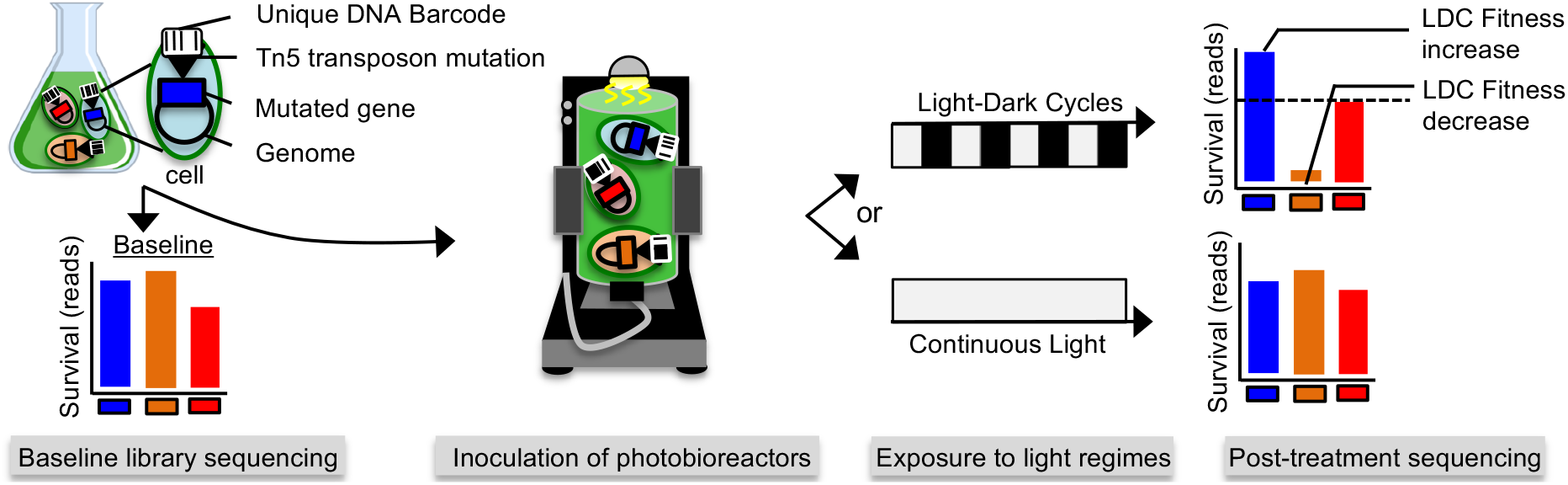
Using RB-TnSeq to assess genetic contributions to fitness in LDC. A dense pooled mutant library containing unique known bar-code identification sequences linked to each individual gene insertion was used to estimate the fitness of each loss-of-function mutant grown under alternating light-dark conditions. The library was grown under continuous light and sampled to determine the baseline abundance of each strain in the population prior to transfer to photobioreactors which were then either exposed to continuous light or alternating 12-h light - 12-h dark regimes. Bar-code quantification of the library population in both conditions was normalized to the baseline and compared against each other to estimate the fitness consequences of loss of function of each gene specific to growth in LDC.

After filtering the data as described in the Materials and Methods section we were able to analyze insertions in 1,872 of the 2,005 non-essential genes (24) in the genome. Each fitness score that passed our statistical threshold for confidence (FDR <1%) was classified as having a “strong” (>1.0 or <-1.0), “moderate” (between 1.0 and 0.5 or −1.0 and −0.5), or “negligible / minor” (between 0 and ±0.5) effect on fitness in LDC. This analysis revealed 90 loci whose disruption caused strong changes to LDC fitness (41 negatively and 49 positively) and an additional 130 loci that had moderate effects (Fig. 2*A*). A volcano plot showing the data with selected genes highlighted in this study is displayed in Fig. 2*B*. Because our initial goal was to understand the suite of genes required to survive the night, we focused first on those mutants whose loss specifically debilitates growth in LDC.

**Fig 2.**
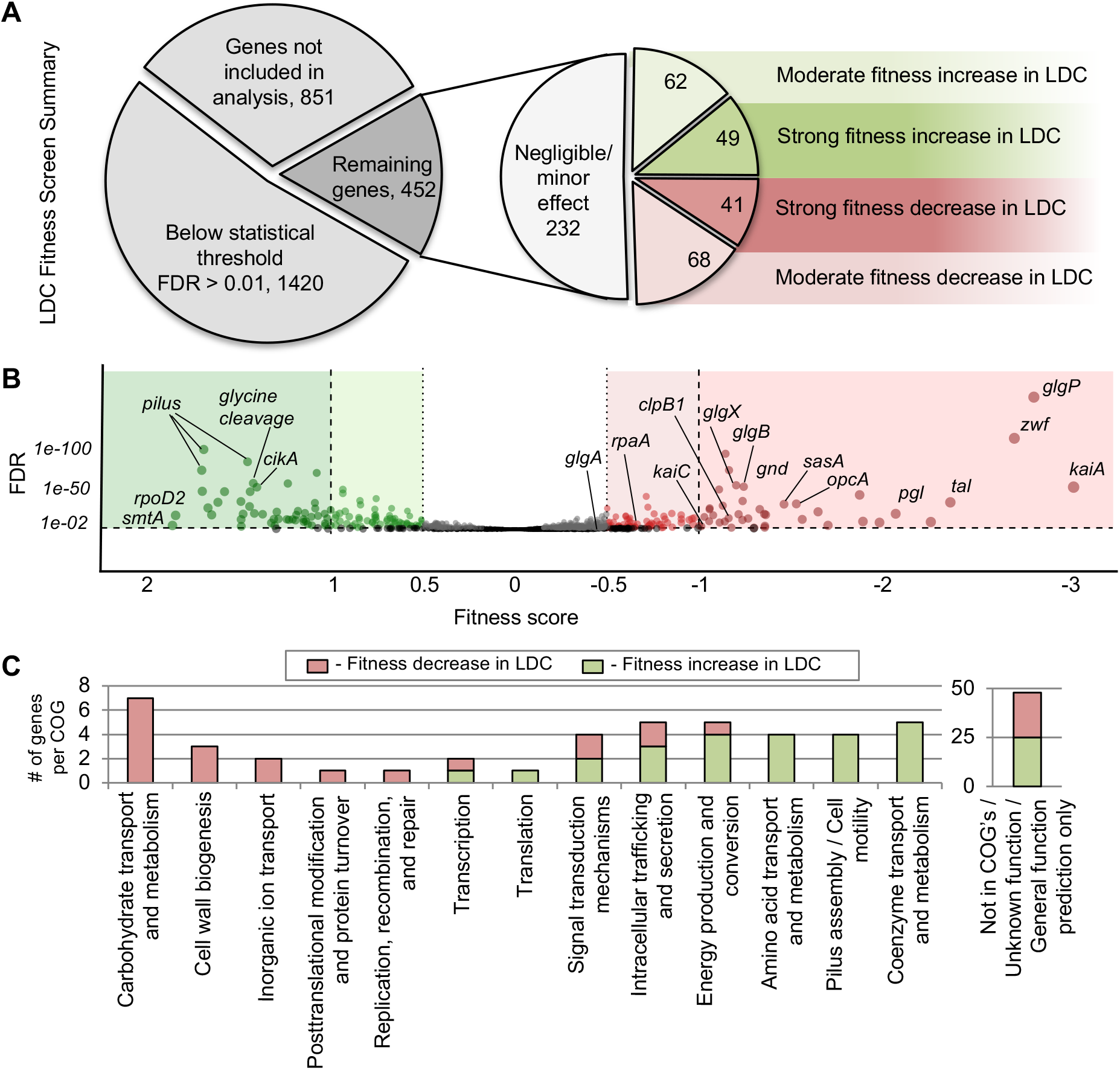
LDC Rb-TnSeq screen results breakdown. *(A)* Of the 2723 genes in the genome, 1872 genes were analyzed in the library population. Of those, data from mutants with loss-of-function of 1420 genes fell below our false discovery threshold. Of the remaining 452 genes, 220 had moderate to strong fitness effects. *(B)* A volcano plot highlighting genes of interest. Red-shaded region indicates an estimated fitness decrease in LDC and the green-shaded region indicates a fitness increase. *(C)* Functional category composition of genes that gave strong fitness scores. Categories based on COG classification or GO ID. Green corresponds to strong fitness increases; Red corresponds to strong fitness decreases.

### Metabolic pathways important for LDC growth

The functional composition of genes found to be either detrimental or favorable to fitness is indicative of the metabolic processes that are important for growth in LDC (Fig. 2*C*). Genes identified as necessary in LDC encode enzymes that participate in carbohydrate and central carbon metabolism, protein turnover and chaperones, CO_2_ fixation, and membrane biogenesis. Strikingly, all genes of the OPPP, *zwf, opcA, pgl*, and *gnd*, have strong fitness effects (Fig. 3*A*). In addition, *tal*, the only non-essential gene of the reductive phase, is also essential for LDC growth. It is known that *tal* mRNA peaks at dusk and that its product acts in a reaction that is shared between the oxidative and reductive phases of the pentose phosphate pathway, linking it to glycolysis; the *tal* mutant’s sensitivity to LDC supports the hypothesis that Tal is a less recognized, but important, component of the OPPP reactions. Furthermore, all OPPP genes have peak expression at dusk (27).

**Fig. 3.**
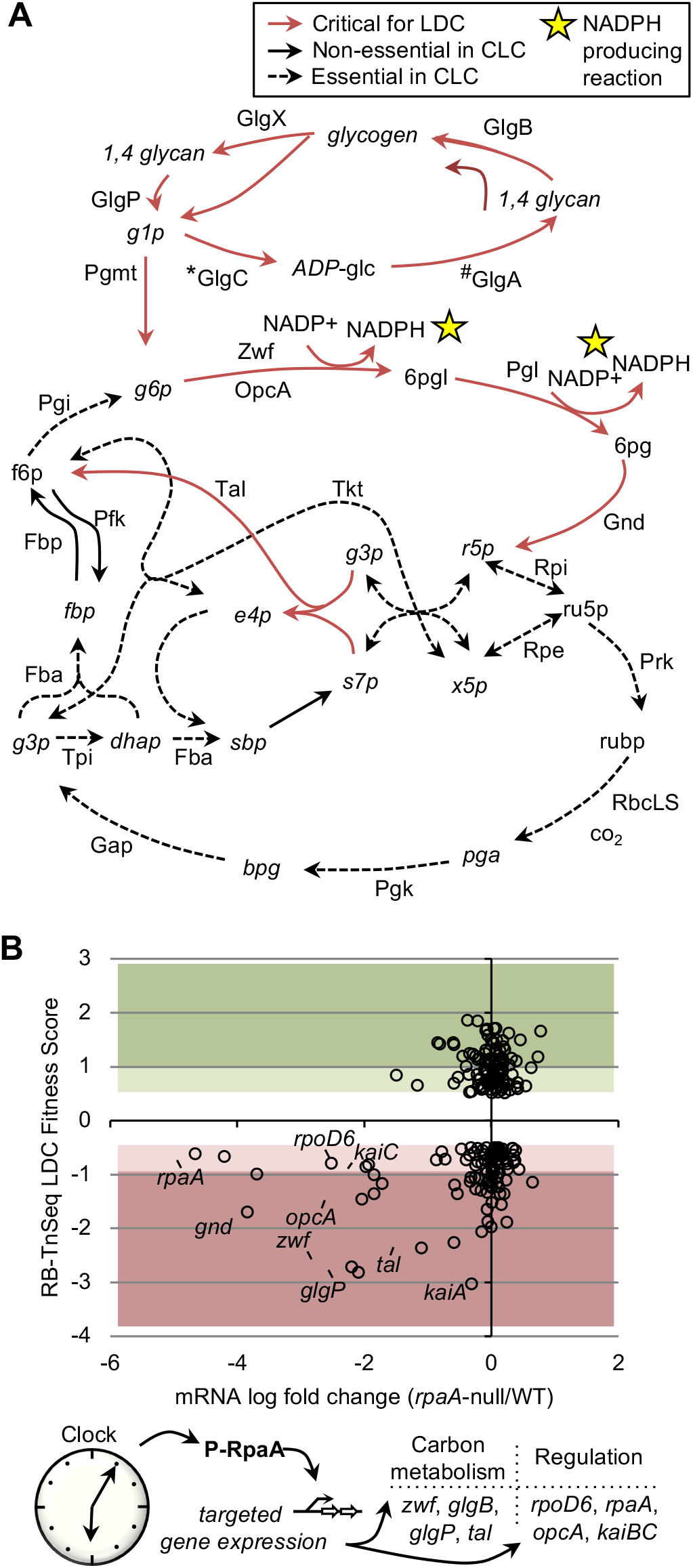
Connection between essential night-time metabolism and RpaA regulation. *(A)* A metabolic map showing the reactions (red) that are controlled by genes that are essential in LDC. Reactions that are non-essential (solid arrows) or essential (dashed arrows) in CLC are are also indicated. Stars mark NADPH-generating reactions. * gene was not revealed in the screen but is known to be required in LDC; # minor LDC decrease measured, likely due to fitness contribution in CLC which affects calculation *(B)* LDC-sensitive mutant population enriched for genes known to be positively regulated by RpaA.

Disruptions in genes that encode enzymes responsible for glycogen metabolism previously have been shown to cause LDC sensitivity (21). Our screen clearly identified the genes for catabolic glycogen metabolism *glgX* and *glgP* as important LDC fitness factors (Fig. 2*B*). However, the importance of glycogen metabolism in LDC fitness is complicated by evidence that the anabolic reactions of *glgA* and *glgC* contribute to fitness in CLC as well, due to the utilization of glycogen as a sink for excess photosynthetically derived electrons (28, 29). Thus, for genes critical for glycogen anabolism, the differential LDC fitness score is not as striking as might be expected from phenotypic assays of those mutants. The *glgA* mutants were classified as having a minor LDC defect, and insufficient reads for *glgC* removed this locus from our analysis pipeline, leading to a classification of “not determined”. Despite this complication, mutations in the gene *glgB*, whose product catalyzes the last non-essential step in glycogen synthesis, caused a strong decrease in LDC fitness, supporting the conclusion that both glycogen synthesis and breakdown are critical aspects of metabolism in LDC conditions. Together, the reactions facilitated by these enzymes provide a major source of reductant (NADPH) in the dark and facilitate the flow of glucose compounds via the breakdown of stored glycogen into downstream metabolites.

Genes involved in DNA repair (such as the transcription-repair coupling factor *mdf*) and chaperone systems *(clpB1)* are also important for light-dark growth. ClpB1 is known to contribute to thermotolerance stress and is induced upon dark-light transitions in cyanobacteria (30, 31). The importance of these genes for surviving light-dark cycles suggests a role in managing light-induced stress that the transition from dark to light may impose. Moreover, mutation of any of the three genes that encode enzymes of the glycine cleavage system confer increased LDC fitness. The glycine decarboxylase complex (GDC) is linked to photorespiratory mechanisms that respond to high-light stress (32, 33). This result further supports our finding that variations in the fitness of metabolism genes between CLC and LDC are linked to pathways that mitigate light stress.

In addition to underscoring the importance of the OPPP, glycogen metabolism, and repair mechanisms in LDC, we also found a strong link between these metabolic pathways and circadian clock management. Many of the genes essential for growth in LDC are regulated by the clock output protein RpaA. For instance, the circadian clock gene *kaiC* is itself regulated by RpaA and its expression is reduced by over 50% in an *rpaA*-null background (12, 14). This attribute is also shared by the core glycogen metabolism and pentose phosphate pathway genes *glgP, zwf, opcA, gnd*, and *tal*, and genes involved with metal ion homeostasis *(smtA)* and cobalamin biosynthesis *(cobL)*, all of which show significantly reduced transcripts (<-1.0 log2(Δ*rpaA*/WT, wild type) in the absence of the RpaA (14). These core glycogen metabolism and pentose phosphate pathway genes, with the exception of *tal* and *gnd*, are also direct targets of the transcription factor. This finding is a common trend in mutants that show strong fitness decreases in LDC, as ~20% (9/41) have dramatic reductions in mRNA levels in an *rpaA*-null strain. This trend is not the case for any of the genes identified as mutants that confer increased fitness (Fig 3*B*). These findings support a growing body of evidence that the ability of *S. elongatus* to initiate a night-time transcriptional program, and particularly to activate the OPPP pathway genes, is critical for survival through darkness (15, 16).

### Fitness contributions of circadian clock constituents

The mechanism of circadian timekeeping and regulation in cyanobacteria is well understood at the molecular level (17, 34). The three core clock proteins, KaiA, KaiB, and KaiC orchestrate circadian oscillation in *S. elongatus* (35), whereby KaiC undergoes a daily rhythm of phosphorylation and dephosphorylation that provides a time indicator to the cell (36, 37). During the day period, KaiA stimulates KaiC autophosphorylation, and, during the night, KaiB opposes KaiA’s stimulatory activity by sequestering an inactive form of KaiA (38), leading to KaiC autodephosphorylation. Two histidine kinase proteins, SasA and CikA, engage with the oscillator at different phases of the cycle. Binding to KaiC stimulates SasA to phosphorylate RpaA, with interaction increasing as KaiC phosphorylation increases. Later, binding of CikA to the KaiC-KaiB complex stimulates CikA phosphatase activity to dephosphorylate RpaA. Thus, activated CikA opposes the action of SasA. This progression causes phosphorylated RpaA to accumulate during the day, peaking at the day-night transition, and activates the night-time circadian program.

Mutations that caused the most serious defect specifically in LDC were mapped to the gene that encodes the clock oscillator component KaiA (Fig. 2*B*). This outcome was unexpected because the known roles of KaiA are to stimulate phosphorylation of KaiC and to enable normal levels of KaiC and KaiB expression (38, 39). Thus, a KaiA-less strain, in which KaiC levels are low and hypophosphorylated (39), was expected to behave like a *kaiC* null. A *kaiC* null mutant has no notable defect in LDC when grown in pure culture (1), and even a *kaiABC* triple deletion, missing KaiA as well as KaiB and KaiC, grows well in LDC (18).

Nonetheless, a *kaiC* null mutant is known to be outcompeted by WT under LDC conditions in mixed culture (1, 2), which is the situation in this population-based screen. Mutations in *kaiC* returned negative fitness scores in the current screen, as did bar codes associated with *rpaA* and sasA, two clock-related genes that are known to be important for growth in LDC (18, 20). RpaA is important for redox balancing, metabolic stability, and carbon partitioning during the night, and an *rpaA* mutant is severely impaired in LDC (15–16, 20). While it is identified in the current screen as LDC sensitive, inactivation of *rpaA* also confers a moderate fitness decrease under CLC (~-1.0), thus diminishing the differential LDC-specific fitness calculation.

Disruptions to other genes related to circadian clock output scored as beneficial to growth in LDC, increasing the fitness of the strains against the members of the population that carry an unmutated copy. For instance, mutations within the gene that encodes CikA showed improved fitness, along with disruptions to the sigma factor encoded by *rpoD2* (40). CikA has several roles that affect the clock, including synchronization with the environment and maintenance of period; additionally, CikA is stimulated to dephosphorylate RpaA by association with the oscillator at night (13, 41). Thus, a phenotype for *cikA* mutants that is opposite to that of *sasA* mutants forms a consistent pattern.

From our previous understanding of the system we would predict that, in a *kaiA* null, SasA kinase would still be activated by KaiC, albeit to a lesser degree than when KaiC phosphorylation levels have been elevated by KaiA. However, CikA phosphatase would not be activated: CikA binds not to KaiC, but to a ring of KaiB that engages hyperphosphorylated KaiC, a state that is not achieved in the absence of KaiA. Therefore, we would expect that RpaA would be constitutively phosphorylated in a *kaiA* null mutant and that the cell would be fixed in the night-time program, where it would be insensitive to LDC. We further investigated the *kaiA-* mutant phenotype under LDC to test the validity of the RB-TnSeq results.

#### Growth attenuation and rescue of a kaiA mutant

The strong negative scores of bar codes for *kaiA* insertion mutants pointed to a severe growth defect in LDC. In order to vet this finding we tested the phenotype of an insertional knockout of the *kaiA* gene *(kaiA^ins^)*, similar to the strains in the RB-TnSeq library. Assays in CLC vs LDC verified the LDC-specific defect in the mutant (Fig. 4*A*). Because neither the *kaiABC* nor *kaiC* deletion mutant has an LDC defect in pure culture, the LDC phenotype seemed to be specific for situations in which KaiC and KaiB are present without KaiA. The *kaiA* coding region has a known negative element located within its C-terminal coding region that negatively regulates expression of the downstream *kaiBC* operon (42, 43) *(kaiA^ins^*), keeping KaiBC levels low. We reasoned that the loss of KaiA would be exacerbated if expression of KaiBC increased. To test this hypothesis, we measured the growth characteristics of a *kaiA* mutant in which the coding region of *kaiA*, including the negative element, has been removed and replaced with a drug marker *(kaiA^del^*) (39). This strain, which is known to have normal or slightly elevated levels of KaiB and KaiC, exhibited even more dramatic sensitivity to LDC and supported the hypothesis that an unbalanced clock output signal underlies the LDC sensitivity in the *kaiA* mutant.

**Fig. 4.**
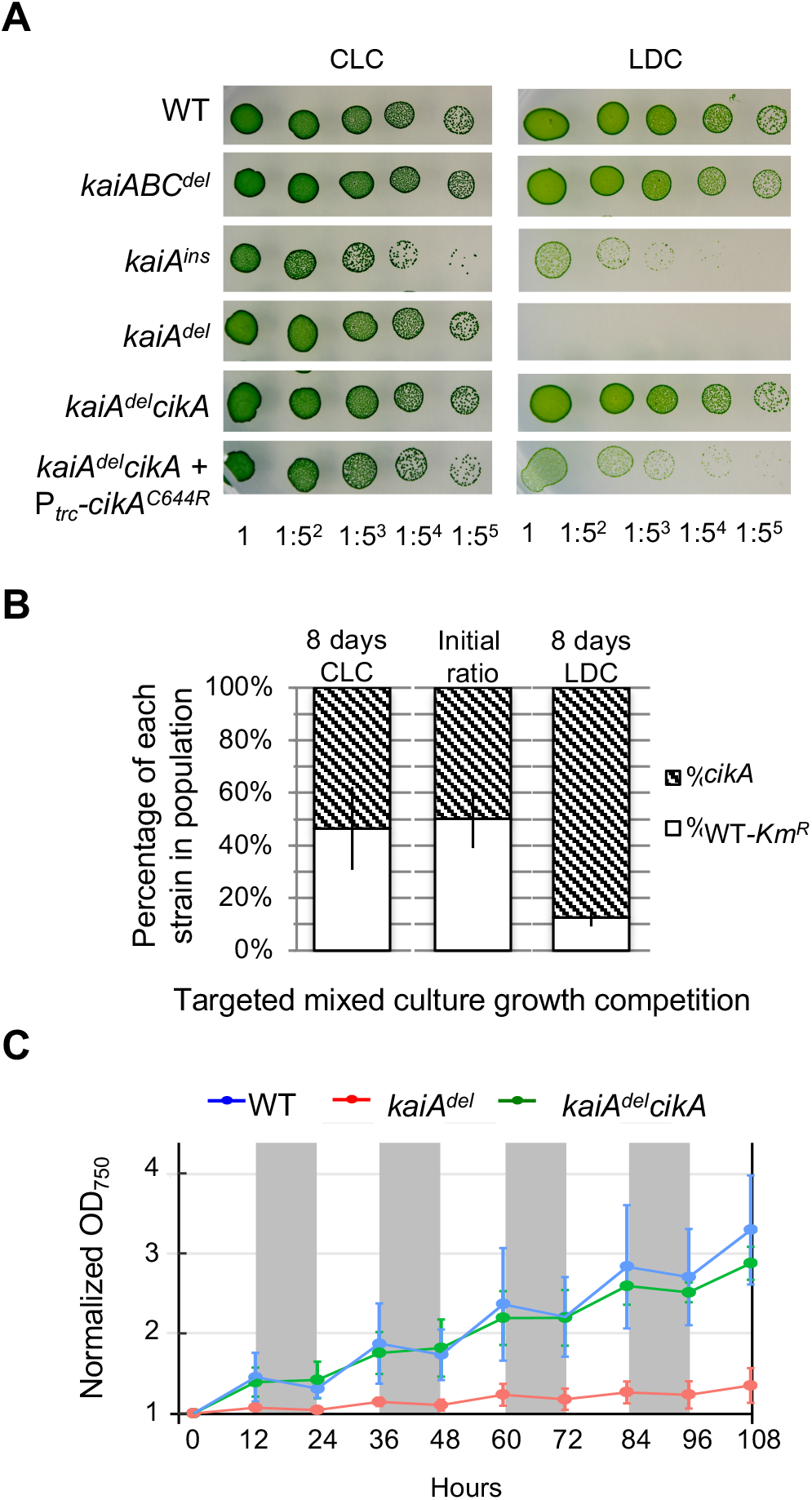
*kaiA* mutant growth and cikA-based intervention in LDC. *(A)* Dilution spot-plate growth of strains in CLC and LDC. A representation of the genotype of each strain is portrayed in SI Appendix, Fig. S1. *(B) cikA* mutant fitness increase in competition with WT in LDC and CLC. *(C)* Growth of liquid cultures in replicate photobioreactors in LDC. Vertical shaded areas represent dark conditions. Data points = mean (SD); n = 2.

Knowing that rpaA-null mutants, which cannot activate the OPPP genes, have similarly severe LDC phenotypes, we hypothesized that the kaiA-less strains have insufficient phosphorylation of RpaA to turn on those genes. We reasoned that additionally interrupting the *cikA* gene, whose product dephosphorylates RpaA in a clock-dependent manner, would help to conserve RpaA phosphorylation in the *kaiA^del^* strain and increase LDC fitness. Indeed, mutations that disrupt *cikA* were identified from the population screen as enhancing fitness (Fig. *2B*), a finding we confirmed in a head-to-head competitive growth assay in a mixed culture with WT (Fig. 4*B*). Complementation of this double mutant with a *cikA^WT^* gene, or the *kaiA^del^* strain with *kaiA^WT^* gene, expressed from a neutral site in the genome, resulted in a reversal of the respective phenotypes (**SI Appendix, Fig. S1**). The LDC growth defects in *kaiA^del^* and improvement of the *kaiA^del^cikA* strain were reproducible when grown in liquid cultures in the same photobioreactors used to grow the library for the LDC screen (Fig. 4*C*). These results are consistent with a model in which, in the absence of KaiA, CikA is constitutively activated as a phosphatase to dephosphorylate RpaA.

#### Arrhythmic and low *kaiBC* expression correlates with LDC phenotype

Hallmarks of RpaA deficiency or dephosphorylation are low and arrhythmic expression from the *kaiBC* locus under CLC, as visualized using a luciferase-based reporter (P_*kaiBC*_::*luc*) (12). We used this assay to determine whether the severity of defect in LDC growth in mutants correlates with level of expression from this promoter as a proxy for RpaA phosphorylation levels. Consistent with the hypothesis, the *kaiA^ins^* mutant has low constitutive levels of luciferase activity, tracking near the troughs of WT circadian expression, the *kaiA* deletion mutant has almost undetectable expression, and the constitutive expression from the *kaiABC^del^* strain tracks near peak WT levels (Fig. 5). Moreover, the *kaiA^del^cikA* strain shows expression levels much higher than the *kaiA^del^* strain, approaching those observed in the *kaiABC^del^* mutant and around the midline of WT oscillation. Intermediate levels of growth in LDC between the *kaiA^del^* and *kaiA^del^cikA* strains were observed when a *cikA* variant (C644R) that is predicted to have reduced KaiB binding, and thus less phosphatase activation, was expressed in *trans* in the *kaiA^del^cikA* strain (11). Growth of this strain also displayed intermediate LDC sensitivity (Fig. 4*A* and **SI Appendix, Fig. S1**). These data support a correlation between expression from the P_*kaiBC*_::*luc* reporter strain, likely reflecting RpaA phosphorylation levels, and growth in LDC.

**Fig. 5.**
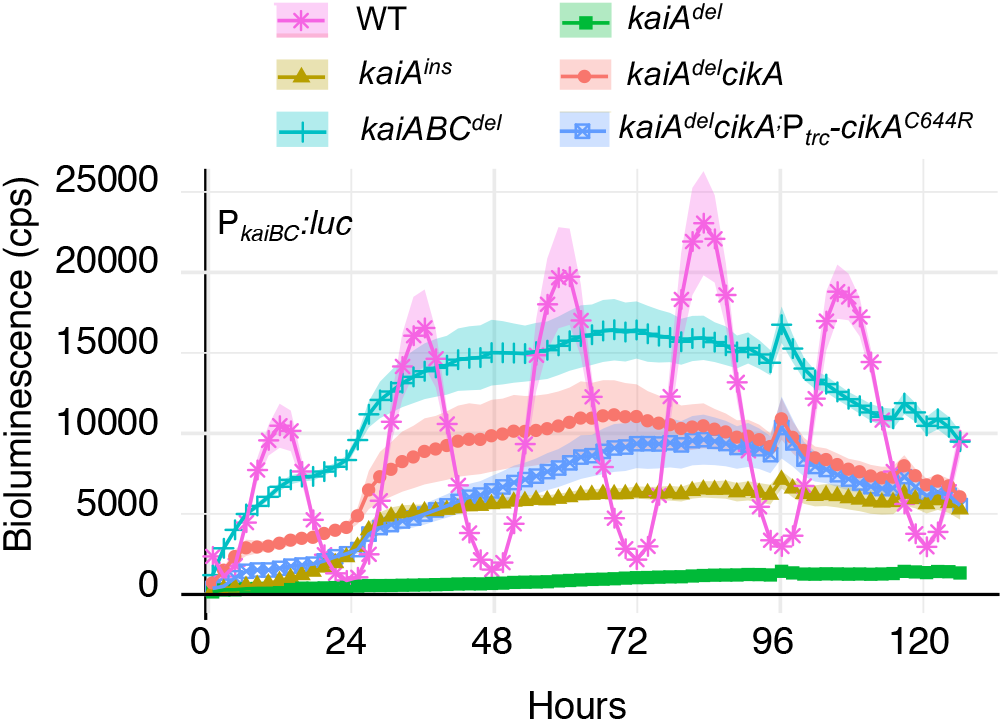
Bioluminescence levels from *kaiBC* reporter in mutant backgrounds. The intensity of *P_kai_B_C_::luc* output signal correlates with LDC sensitivity. Rhythms were measured in CLC conditions driven by the *kaiBC* promoter after exposure to 72 h of entraining LDC that includes a low-light intensity that is permissive for the mutants. Shaded areas indicate the SEM of 4 biological replicates.

#### Low RpaA phosphorylation in kaiA-null cells as the cause of decreased fitness

We next directly assayed the RpaA phosphorylation levels in this group of mutants by using immunoblots and Phos-tag reagent to separate phosphorylated (P-RpaA) from unphosphorylated RpaA. As predicted, P-RpaA was low in the *kaiA^ins^* mutant and almost undetectable in the *kaiA^del^* strain (Fig. 6*A*). Adding an ectopic WT *kaiA* allele, or disrupting *cikA*, in the *kaiA^del^* mutant restored RpaA phosphorylation. Thus, even though adequate KaiC is present to engage SasA to phosphorylate RpaA (Fig. 6*B*), the absence of KaiA is sufficient to suppress the accumulation of phosphorylated RpaA. These results support the conclusion that the rescue of the *kaiA* growth defect by disrupting *cikA* is due to the elimination of CikA phosphatase activity.

**Fig. 6.**
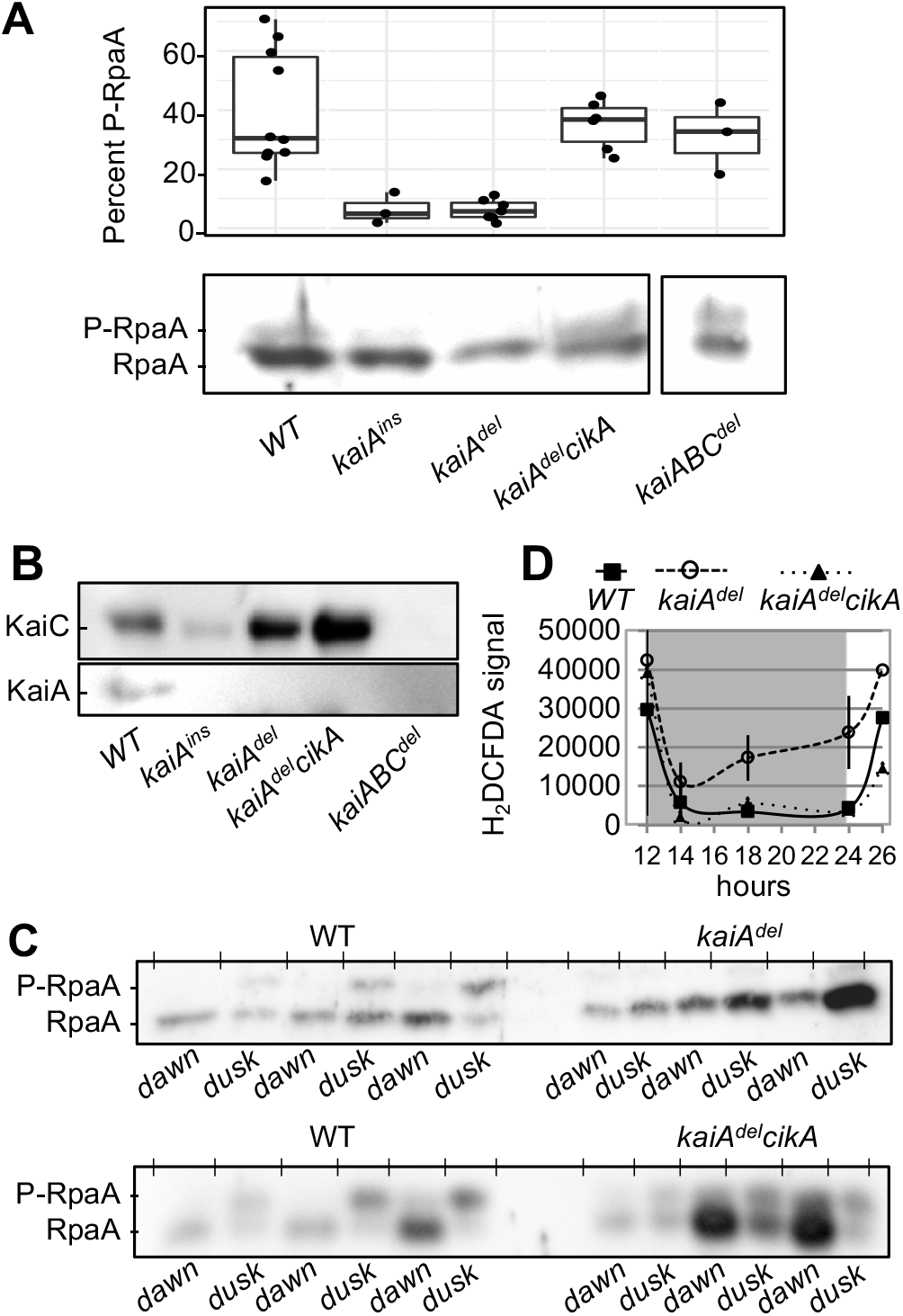
RpaA status correlates with growth success in LDC. *(A)* RpaA phosphorylation at dusk (after 12 h light). Box plot above the representative Phos-tag immunoblot shows values for replicate densitometry analysis (n ≥ 4) of P-RpaA/total RpaA. *(B)* Immunoblot shows levels of KaiC from cells taken at dusk. C. Phos-tag immunoblot shows RpaA phosphorylation across the light-dark transitions with samples taken just prior to the lights turning on (dawn) or off (dusk). *(D)* H_2_DCFDA fluorescence over a 12-h dark period during an LDC, indicating total cellular ROS in WT, *kaiA^del^*, and *kaiA^dol^cikA.* The shaded area indicates the period of darkness following a 12-h light period. P-RpaA denotes phosphorylated protein.

Because the model suggests that turning on RpaA-dependent genes at dusk is critical for survival in LDC, we measured P-RpaA specifically at light-to-dark and dark-to-light transitions in *kaiA^del^* and *kaiA^del^cikA* over three diel cycles. WT has a marked pattern of high P-RpaA at dusk and low levels at dawn (Fig. 6*C*). The *kaiA^del^* strain had almost undetectable levels of P-RpaA, visible only when the overall RpaA signal was very high. The P-RpaA phosphorylation status in the *kaiA^del^cikA* was restored but did not show diel variations, consistent with the arrhythmicity of *kaiA* mutants.

Other physiological similarities between the kaiA-null and rpaA-null strains exist. An *rpaA* mutant accumulates excessive reactive oxygen species (ROS) during the day that it is unable to alleviate during the night (15). Metabolomic analysis points towards a deficiency in reductant, needed to power detoxification reactions for photosynthetically-generated ROS, due to lack of NADPH production in the dark when OPPP genes are not activated. Likewise, levels of ROS in *kaiA^del^*were significantly higher at night than in WT and *kaiA^del^cikA* (Fig. *6D*).

#### Molecular basis of CikA engagement in the absence of KaiA

The proposal that the LDC fitness defects in *kaiA* mutants result from excessive CikA phosphatase activity was derived from genetic data related to known LDC-defective mutants. However, previous studies regarding KaiA activities had not predicted this outcome. Very recent structural data, enabled through the assembly of KaiABC and KaiB-CikA complexes, pointed to a likely mechanism: loss of competition for a binding site on KaiB that KaiA and CikA share (11). Although this hypothesis was appealing, the current model of progression of the Kai oscillator cycle could not predict this outcome. KaiA and CikA bind to a ring of KaiB that forms after KaiC has become phosphorylated at Serine 431 and Threonine 432 (44). Maturation to this state induces intramolecular changes in KaiC that expose the KaiB binding site. In the absence of KaiA, KaiC autophosphorylation is extremely abated as measured both *in vivo* (45) and *in vitro* (1-10%, **SI Appendix, Fig. S5C**), so how can KaiB bind?

We used fluorescence anisotropy to directly determine whether KaiB can engage with KaiC in the absence of KaiA, when KaiC is mainly unphosphorylated (Fig. 7 and **SI Appendix, Fig. S5**). If KaiC-KaiB binding can occur, it could support binding of CikA – expected to be a necessary event for CikA phosphatase activation. This method measures the magnitude of polarization of fluorescence from KaiB tagged with a fluorophore (6-iodoacetamidofluorescein), which increases when KaiB becomes part of a larger complex. KaiC phosphomimetic mutants were used to quantify interaction of KaiB with each of the phosphostates of KaiC. Initially, we hypothesized that the KaiB binding site might become exposed on KaiC in the absence of phosphorylation when cells enter the dark and the ATP/ADP ratio is altered (46). For this reason, experiments were performed at different ATP/ADP ratios. As a control, we confirmed that no significant association of KaiB with CikA occurs in the absence of KaiC (Fig. 7*A*). However, KaiB is able to associate with hypophosphorylated WT KaiC approximately as well as with the phosphomimetic, KaiC-EA, which mimics a form of KaiC (KaiC-pST, where S431 is phosphorylated) that binds KaiB, which then forms a new ring on the N-terminal face of KaiC that captures KaiA and CikA. This binding was similar in both 1 mM ATP (Fig. 7*B*) and 0.5 mM ATP+0.5 mM ADP conditions (Fig. 7*A*), suggesting that darkness (associated with decreased ATP/ADP *in vivo)* is not required to enable KaiB binding to KaiC. Moreover, CikA associated with KaiBC under both conditions. Association of KaiB was minimal with KaiC-AE, a mimic of the daytime state (KaiC-SpT, where T432 is phosphorylated) when KaiA is usually well-associated with the A-loops of KaiC in an interaction that does not require KaiB. However, the presence of CikA enhanced the binding of KaiB to KaiC-AE, presumably due to cooperative formation of a KaiC-KaiB-CikA complex. For the WT KaiC experiments in Fig. 7, 6IAF-KaiB (0.05 μM) is most likely binding to the small percentage of the KaiC population that remains phosphorylated (1-10% = 0.04-0.4 μM). *In vivo*, it is likely that in the absence of KaiA, KaiB likewise binds selectively to a small fraction of the KaiC pool that is phosphorylated.

**Fig. 7.**
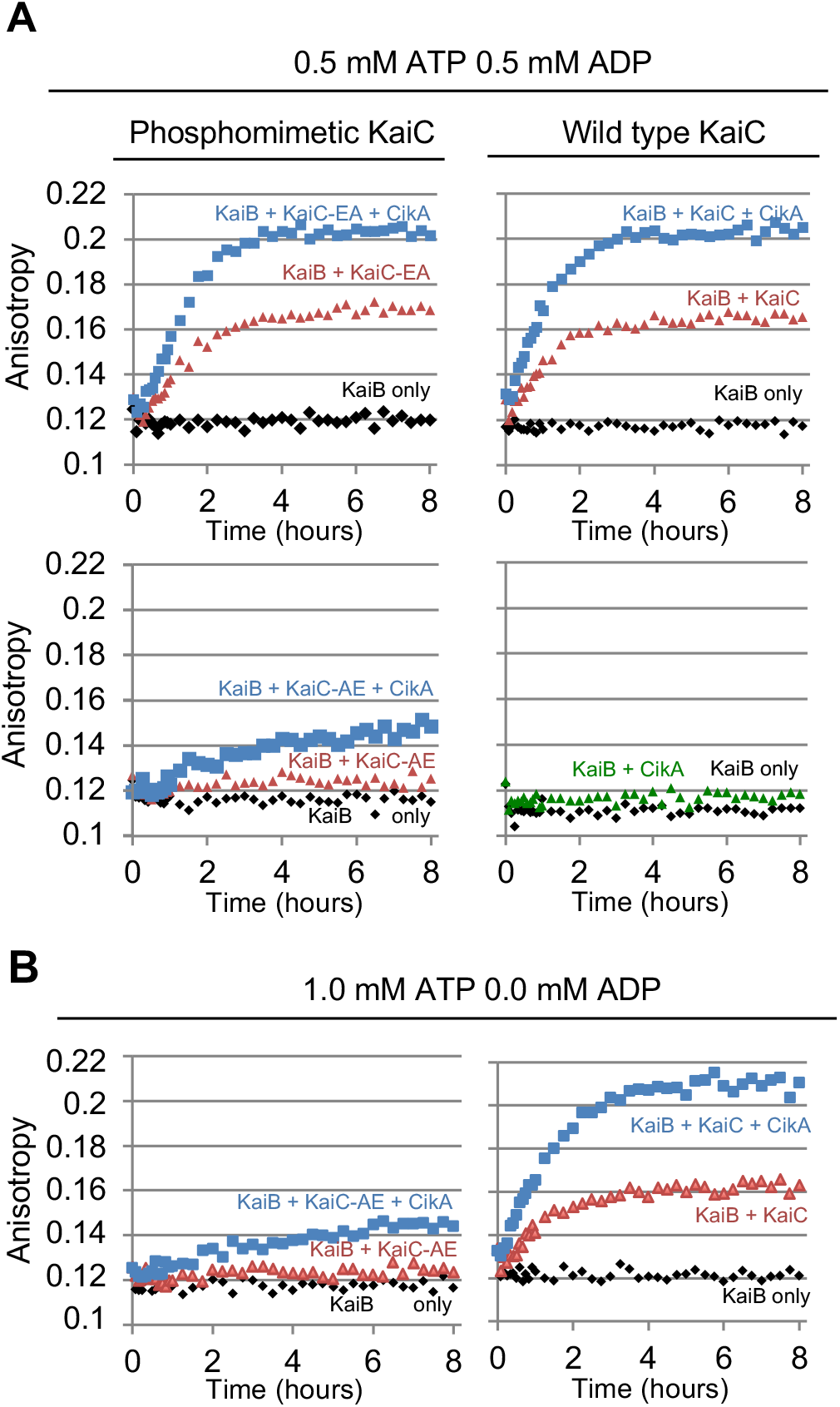
CikA-activating complex formation measured by fluorescence anisotropy. *(A)* Kinetics of 6-IAF-labeled KaiB binding to WT KaiC (upper right) or phosphomimetic KaiCs (KaiC-AE, or KaiC-EA, left) in the absence (red) or presence (blue) of CikA, with 0.5 mM ATP + 0.5 mM ADP. Controls are shown for KaiB-only (black) and for KaiB and CikA without KaiC (green, lower right). *(B)* Binding of KaiB to WT KaiC (right) or KaiC-AE (left) in the absence and presence of CikA with 1.0 mM ATP, using the same color codes as for (A).

These data generally match the patterns expected for the current oscillator model; however, they also demonstrate that the KaiC-KaiB-CikA complex can form in the absence of KaiA, when the KaiC pool is mostly hypophosphorylated, and that the ATP ratio (mimicking day and night) has little effect on this association. Furthermore, they support the competitive binding of KaiA and CikA to the KaiB ring. These observations are consistent with the proposal that the low survival rate of *kaiA-null* mutants is due to over-stimulation of CikA phosphatase activity, stripping the cell of P-RpaA and disabling expression of the critical night-time reductant-producing OPPP.

## Discussion

Recent research and interest in growth of cyanobacteria in LDC conditions (15, 16, 20, 22, 47–49) provides a rich dataset to inform engineering strategies, to model metabolic flux predictions, and to elucidate the intersection of the clock and cell physiology. While previous studies have identified a handful of genes important for growth in LDC, the conditional defect of these mutants was determined from targeted studies that left large unknowns in the pathways important for LDC. This study used an unbiased quantitative method to comprehensively query the *S. elongatus* genome for the full set of genes that are specifically necessary to survive the night. This approach eliminated the requirement to test individual mutants for a scorable phenotype under diel conditions and allowed for the discovery of genes that would not be predicted to have such a phenotype. In addition to identifying genes of pathways previously unknown to be involved in LDC, this screen also strongly reinforced two emerging stories from recent research: the breakdown of glycogen and flux through the OPPP, the major source of reductant under non-photosynthetic conditions, is essential (15, 16); and, the role of the clock is repressive, rather than enabling (12), as its complete elimination has little consequence in LDC, but its constitutive output signaling is harmful under these conditions because of its negative effect on the OPPP genes.

A boon of the Rb-TnSeq data is in how it reveals non-intuitive insights into the regulatory measures cells need for growth in specific conditions. One such discovery is that a *kaiA* mutant is severely LDC sensitive. Investigation of the mechanism that underlies this phenotype led to increased understanding of the function of the Kai oscillator, a nanomachine whose structural interactions have been revealed with an unusual degree of clarity (11, 50). KaiA is conventionally thought to have a specific daytime role in stimulating KaiC autophosphorylation – an activity that promotes SasA stimulation and brings about the conformational change in KaiC that enables KaiB engagement (Fig. *8A*). This latter step ushers in the KaiC dephosphorylation phase, during which KaiA is a passive, inactive player. Results from this study provide a new perspective of KaiA in its KaiB-bound form as regulating the night-time clock output by its competition with CikA for KaiB-binding (Fig. 8*B*). A cell without KaiA leaves the major transcription factor RpaA hypophosphorylated for two reasons. First, without KaiC undergoing its daytime phosphorylation cycle, SasA engagement with the oscillator is diminished, reducing its kinase activity acting on RpaA. Secondly, without KaiA present as a competitor, the opportunities for CikA binding on KaiB-KaiC complexes increase, leading to hyperactive phosphatase activity and ultimately eliminating any chance for phosphorylated RpaA to accumulate. The result is a severe decrease in fitness in LDC. However, cells without KaiA and CikA (and even without KaiABC) have ample P-RpaA, indicating that other pathways contribute to the phosphorylation of RpaA. This fact emphasizes the counterintuitive point that activation of CikA phosphatase in the night-time complex, rather than activation of SasA kinase, is the key signaling state of the cyanobacterial clock (12).

**Fig. 8.**
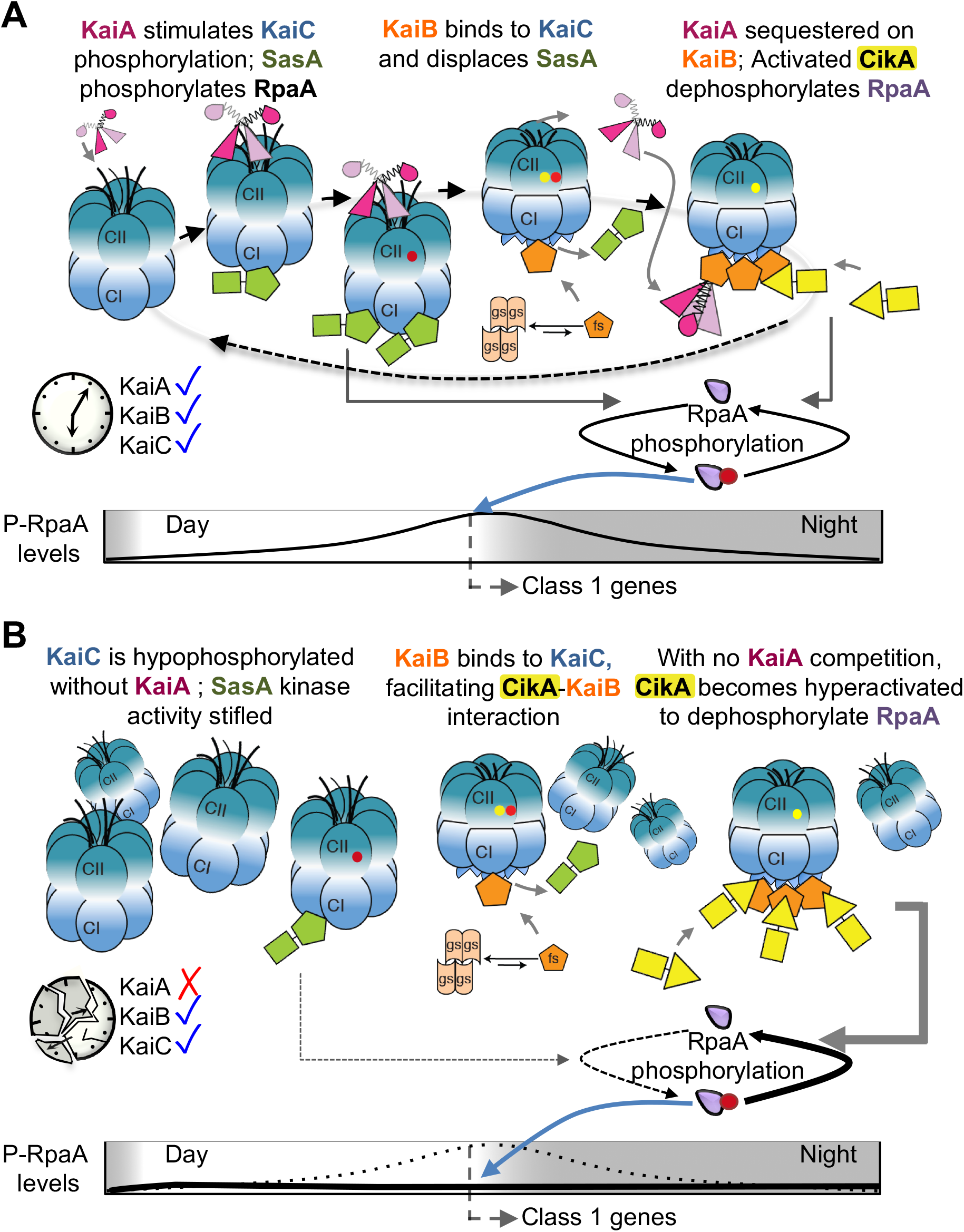
Model for LDC sensitivity due to circadian clock disfunction. *(A)* Conventionally, the role of KaiA is primarily to bind to the A-loops of the CII domain of KaiC, stimulating its autophosphorylation and consequently increasing the activity of the kinase SasA; this activity in turn results in accumulation of phosphorylated RpaA throughout the day. P-RpaA initiates the expression of circadian-controlled night-time class 1 genes at dusk. Yellow and red dots on KaiC represent phosphorylation at S431 and T432, respectively. A red dot also represents phosphorylation on RpaA. The rare fold-switch of KaiB is represented by different orange shapes. The major conformational change of KaiA bound to the CII A-loops versus the KaiB ring and the exposure of the KaiB binding site on the CI ring of KaiC are also depicted. *(B)* Without KaiA, KaiC phosphorylation is suppressed and, thus, SasA kinase activity is diminished, reducing the pools of SasA-mediated P-RpaA. A small fraction of the KaiC pool that is phosphorylated even in the absence of KaiA can bind KaiB. CikA has access to these KaiC-bound KaiB proteins that are usually occupied by KaiA, and becomes hyperactive; excessive CikA phosphatase activity extinguishes intracellular pools of P-RpaA and eliminates the cell’s ability to express genes needed for survival in LDC. Figure modified from (11). The dotted line in the panel B graph represents the WT levels of P-RpaA from panel A.

The screen revealed expected mutants as well. Although a *kaiC* mutant does not have a notable LDC defect in pure culture, it is known to do poorly in LDC when grown in a population with other cells that have a functioning circadian rhythm, as is the case in the pooled growth scenario used here, illustrating soft selection for circadian rhythms in cyanobacteria (1). As expected, *kaiC* was deficient under LDC, but not as severely as many other mutants. One of the most severely LDC-defective mutants we know of is *rpaA*, which dies within a few hours of dark exposure and requires special handling in the lab. Strains with disruptions in *rpaA* did score as negative, but only to a moderate degree (−0.7). This discrepancy between severe known phenotype and moderate score can be attributed to the fact that *rpaA* mutants were also less fit than the population in CLC, decreasing its magnitude of relative fitness in LDC, similar to the situation for mutants defective in glycogen anabolic processes.

Importantly, the screen also identified an equivalent list of genes whose loss improves growth in LDC. Although not the target of the current study, these genes have great potential for providing new insights into *S. elongatus* diurnal physiology and perhaps improved genetic backgrounds for metabolic engineering. In contrast to many of the mutants that negatively affect LDC growth, the list of top positive mutants includes many genes with less transparent roles in LDC conditions, including some that carry no functional annotation. The phenotype may be attributable to a specific pathway or may result from the additive effects of many unrelated genes whose expression is altered in a regulon indirectly. Some mutants may outcompete others in the population because they have alleviated photochemical intermediate buildup or lessened the uptake of harmful metabolites in the medium. The scant information available regarding these pathways makes it difficult to predict why their value would be different in CLC vs. LDC.

In summary, this study has improved our understanding of the mechanism of core oscillator protein interactions and generated a comprehensive map of genetic pathways needed for survival in LDC in *S. elongatus*. We have shown that KaiA, originally thought to only regulate KaiC autophosphorylation, also modulates CikA-mediated clock output. More broadly, the use of RB-TnSeq on a dense transposon library in this study has identified the genes in the genome necessary for growth in alternating light-dark conditions, established a firm connection between circadian clock regulation and the metabolic pathways critical for fitness in more natural environments, and contributed to a more complete picture of the finely balanced mechanisms that underlie circadian clock output.

## Materials and Methods

### Bacterial strains and culture conditions

All cultures were constructed using WT *S. elongatus* PCC 7942 stored in our laboratory as AMC06. Cultures (**Table S1**) were grown at 30 °C using either BG-11 liquid or solid medium with antibiotics as needed for selection at standard concentrations: 20 μg/mL for kanamycin (Km) and gentamicin (Gm), 22 μg/mL for chloramphenicol (Cm), and 10 μg/mL each for spectinomycin and streptomycin (SpSm) (51). Liquid cultures were either cultivated in 100 mL volumes in 250 mL flasks shaken at ~150 rpm on an orbital shaker or in 400 mL volume in top-lit bioreactors (Phenometrics Inc. ePBR photobioreactors version 1.1) mixed via filtered ambient-air bubble agitation with a flow rate of 0.1 mL per min at 30 °C. Viable cell plating for LDC-sensitivity testing and quantification of cellular ROS via 2’,7’-dichlorodihydrofluorescein diacetate (H_2_DCFDA) were performed as previously described (15). Spot-plate growth was quantified by converting the plate image to 8-bit and performing densitometric analyses using National Institutes of Health ImageJ software (52). Transformation was performed following standard protocols (53). Strains were checked periodically for contamination using BG-11 Omni plates (BG-11 supplemented with 0.04% glucose and 5% LB) (53). Verification of the disruption of the *kaiA* gene in mutants was performed by PCR and is shown in **SI Appendix, Fig. S1**. For immunoblotting experiments and ROS measurements cultures were grown in flasks. Samples for protein were taken after 3 light-dark cycles at specified time points immediately before the lights turned on and turned off.

### One-on-one competition experiments

The relative fitness of *cikA* null and *cikA+* strains was assessed in growth competition experiments that leveraged different antibiotic resistance markers in the strains. The *cikA* mutant carries a Gm-resistance marker inserted in the *cikA* locus. AMC06 was transformed with pAM1579 to generate a Km-resistant strain that carries the drug cassette in neutral site II (designated as “WT-Km^R^” for this experiment). The axenic cultures were grown to OD_750_ ~ 0.4, washed three times with BG-11 via centrifugation at 4,696 G for 10 min to remove antibiotics, and diluted with BG-11 to an OD_750_ of 0.015. Cultures were then mixed in a 1:1 ratio and grown without antibiotics for 8 days under ~100 μmol photos · m^2^ · s^−1^ light. To assay survival of each genotype at 0, 3, and 8 days, mixed cultures were diluted using a 1:10 dilution series and plated in 20 μL spots in duplicate onto ~2 mL of BG-11 agar with either Gm or Km poured in the wells of a 24-well microplate. CFU’s of each strain were calculated from the dilution series across the two antibiotic regimes and were used to determine the percentage of each strain in the mixture population over time. To ensure that the expression of the antibiotic markers did not influence fitness, WT cultures that carried each drug-resistance marker (Km in neutral site II via plasmid pAM1579 or Gm in neutral site III via plasmid pAM5328) were grown together and assayed, and displayed no significant differences (**SI Appendix, Fig. S2**).

### Rb-Tnseq Assay

Samples of the library archived at −80 C were quickly thawed in a 37 C water bath for 2 min and then divided into three flasks of 100 mL BG-11 with Km, and incubated at 30 °C in 30 μmol photos · m^2^ · s^−1^ light for 1 day without shaking. The library culture was then moved to 70 μmol photos · m^2^ · s^−1^ light on an orbital shaker till OD_750_ reached ~0.3. The cultures were then combined, diluted to starting density of OD_750_ 0.025, and 4 replicates of 15 mL were spun down at 4,696 g and frozen at −80 °C as Time 0 samples to determine population baseline. After transfering to bioreactors, CLC bioreactors were set for continuous light at 500 μmol photos · m^2^ · s^−1^ and LDC bioreactors were set to a square-wave cycle consisting of 12 h light - 12 h dark. Bioreactors were set to run in turbidostat mode maintaining the density at OD_750_ = 0.1. Bioreactors were sampled after approximately 6 generations: the CLC reactors after 3 days (three 24-h light periods) and LDC reactors after 4 days (four 12-h light-12-h dark light dark periods). Two of the three biological replicates for each condition were performed concurrently (using 4 bioreactors) while a third was performed later using the same procedure and conditions.

### Bioluminescence monitoring

Flask-grown batch cultures that reached a density of OD_750_ = 0. 3-0.4 were diluted as needed to OD_750_ of 0.3 and 20 μL of each culture was placed on a pad of BG-11 agar containing 10 μL of 100 mM firefly luciferin, in a well of a 96-well dish. Plates were covered with clear tape to prevent drying and holes were poked using a sterile needle to allow air transfer. Cultures were synchronized by incubating the plate under permissive LDC conditions (0 and 30 μmol photos · m^2^ · s^−1^ light) for 3 days, and then returned to constant light (LL) conditions for bioluminescence sampling by a Packard TopCount luminometer (PerkinElmer Life Sciences). Bioluminescence of *P_ka_¡B_C_::luc* firefly luciferase fusion reporter was monitored at 30 °C under CLC as described previously (54). Data were analyzed with the Biological Rhythms Analysis Software System (http://millar.bio.ed.ac.uk/pebrown/brass/brasspage.htm) import and analysis program using Microsoft Excel. Results shown are the average of four biological replicate wells located in the innermost section of the plate, where drying is minimal.

### Protein sample preparation and gel electrophoresis, and Phos-tag acrylamide gels

Protein was isolated from cells and diluted to final loading concentration of 5 μg/well for RpaA analysis and 10 μg/well for KaiC analysis (55). SDS/PAGE was performed according to standard methods with the following exceptions. Phosphorylation of RpaA was detected using 10% SDS-polyacrylamide gels supplemented with Phos-tag ligand (Wako Chemicals USA) at a final concentration of 25 μM and manganese chloride at a final concentration of 50 μM. Gels were incubated once for 10 min in transfer buffer supplemented with 100 mM EDTA, followed by a 10-min incubation in transfer buffer without EDTA before standard wet transfer. Protein extracts and the electrophoretic apparatus were chilled to minimize hydrolysis of heat-labile phospho-Asp. Protein extracts for use in Phos-tag gels were prepared in Tris-buffered saline, and extracts for standard SDS/PAGE were prepared in PBS. RpaA antiserum (a gift from E. O’Shea, Harvard University, Cambridge, MA) was used at a dilution of 1:2,000 and secondary antibody (goat anti-rabbit IgG, Calbiochem Cat # 401315) at 1:5000. KaiC immunoblotting was performed similarly using KaiC antiserum diluted to 1:2000 and secondary antibody diluted to 1:10,000 (Horseradish Peroxidase-labeled goat anti-chicken IgY, Aves Labs, Cat # H-1004) (56). Densitometric analyses were performed using National Institutes of Health ImageJ software (52) (**SI Appendix, Fig. S3**, Dataset S3). Samples were immediately stored at −80 °C, thawed once, and always kept on ice.

### Fitness calculations

Of the 2,723 genes comprising the genome of *S. elongatus* 718 are essential, and mutants in the essential genes are not present in the library (24). To estimate the fitness effects of gene disruptions in LDC relative to CLC, we developed an analysis pipeline of curating the data from the 2,005 nonessential genes, normalizing it, and then analyzing it using linear models (**Dataset S1**). We first counted the number of reads for each sample to use as a normalizing factor between samples. Bar codes are dispersed across the genome, and we removed any bar code falling outside of a gene (24,868 bar codes out of 154,949 total bar codes) or within a gene but not within the middle 80% (27,763/154,949). Based on the bar codes remaining, we removed any gene not represented by at least three bar codes in different positions (114 genes out of 2,075 total). This filtering left us with 102,136 bar codes distributed across 1,961 genes.

For each bar code in each sample we added a pseudocount of one to the number of reads, divided by the total number of reads for the sample as calculated before, and took the log-2 transformation of this sample-normalized number of reads. The experiment involved two different starting pools of strains (called T0), each of which was divided into CLC and LDC samples. To account for different starting percentages of each bar code within the T0 pools, we averaged the log-2 transformed values for a bar code across the four replicate T0 samples for each pool then subtracted these average starting bar-code values from the CLC and LDC values in the respective pools. We also removed any gene without at least 15 T0 reads (across the four replicates and before adding the pseudocount) in each pool (89/1,961), leaving 1,872 genes and 101,258 bar codes.

For each gene, we used maximum likelihood to fit a pair of nested linear mixed effects models to the sample- and read-normalized log-2 transformed counts:

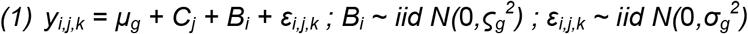

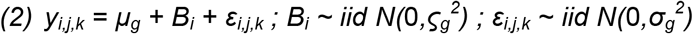

where *y_i,j,k_* is the normalized log-2 value for bar code *i* in gene *g* in condition *j* for sample *k, μ_g_* is the average value for the gene, *C_j_* is the fixed effect of condition *j, B_i_* is a random effect for bar code *i*, and *ε_i,j,k_* is the residual. We identified genes with significant fitness differences between conditions by comparing the difference in the −2*log likelihoods of the models to a chi-square distribution with one degree of freedom, estimating a p-value, accounting for multiple testing by the false-discovery rate method of Benjamini and Hochberg (57), and selecting those gene with adjusted p-values less than 0.01. We took the contrast *C_LDC_ – C_CLC_* to be the estimated LDC-specific fitness effect of knocking out the gene (**Dataset S2**).

Results obtained across biological replicates and across similar screens performed in either batch cultures grown in 250 mL Erlenmeyer shake flasks and on solid agar on Petri plates were consistent (**SI Appendix, Fig. S4, Dataset S4**). Mutants that had estimated fitness scores below a confidence threshold (associated FDR adjusted p-value <0.01) (Dataset S2) were not considered. Mutants with gene disruptions that had an LDC fitness score ≥ 1.0 or ≤ −1.0 with a false discovery rate ≤ 1% were classified having strong fitness phenotype specific to growth in LDC, and those with a fitness score of < 1.0 but ≥ 0.5 and > −1.0 but ≤ −0.5 were classified as causing a moderate fitness phenotype. Fitness scores <0.5 and >-0.5 were categorized as negligent/minor, i.e., little change in abundance over the course of the experiment in LDC vs. CLC.

### Fluorescence Spectroscopy

To monitor the formation of the CikA activating complex, fluorescence anisotropy measurements were performed at 30 °C on an ISS PC1 spectrofluorometer equipped with a three-cuvette sample compartment, at excitation and emission wavelengths of 492 nm and 530 nm, respectively, for samples including 6-iodoacetamidofluorescein (6-IAF) labeled KaiB-FLAG-K251C alone (0.05 μM), and its mixtures with KaiC (4 μM) or FLAG-CikA (4 μM) or KaiC (4 μM) + FLAG-CikA (4 μM). Each sample (400 μL) was either in an ATP buffer [20 mM Tris, 150 mM NaCl, 5 mM MgCl_2_, 1 mM ATP, 0.5 mM EDTA, 0.25 mM TCEP (tris(2-carboxyethyl)phosphine), pH 8.0] or in ATP/ADP buffer [20 mM Tris, 150 mM NaCl, 5 mM MgCl_2_, 0.5 mM ATP, 0.5 mM ADP, 0.5 mM EDTA, 0.25 mM TCEP, pH 8.0]. TCEP was introduced from the FLAG-CikA stock [20 mM Tris, 150 mM NaCl, 5 mM TCEP, pH 8.0]. KaiB stock was in the buffer [20 mM Tris, 150 mM NaCl, pH 8.0] and KaiC stock in the buffer [20 mM Tris, 150 mM NaCl, 5 mM MgCl_2_, 1 mM ATP, 0.5 mM EDTA, pH 8.0].

The KaiC samples included freshly dephosphorylated WT KaiC, and KaiC phosphomimetics KaiC-AE and KaiC-EA. To prepare the freshly dephosphorylated KaiC, WT KaiC at 20 μM in the buffer [20 mM Tris, 150 mM NaCl, 5 mM MgCl_2_, 1 mM ATP, 0.5 mM EDTA, pH 8.0] was incubated at 30 °C for 24 hours before the fluorescence binding assays. SDS-PAGE analysis was performed to determine KaiC and CikA stability and KaiC phosphorylation levels (**SI Appendix, Fig. S5, Dataset S5**).

## Acknowledgments

We thank L. Lowe and J. Tan for technical assistance. This work was supported by research grants (R35GM118290 to SSG; R01GM107521 to AL) and training grants (T32GM007240 to BER and SD) from the National Institutes of Health and the National Science Foundation (MCB1517482 to SAR).

