## Supplementary material for "Genome-wide fitness assessment during diurnal growth reveals an expanded role of the cyanobacterial circadian clock protein KaiA": LDScreen_BioreactorsAnalysis.nb.html (from another version of this preprint: version 1, 2018-03-17)

Light-Dark RB-TnSeq Screen


Code 

- Show All Code
- Hide All Code
- Download Rmd

### Light-Dark RB-TnSeq Screen

Using data from continuous light and light-dark growth conditions in photobioreactors. The time zero from group1 was placed in a flask and divided up into two experimental and one control and the second grouping time0 was divided into one experimental and one control. Each Group of time0’s have quadruplal technical reps.

Tidy business code


```
library(knitr)
opts_chunk$set(tidy.opts=list(width.cutoff=60),tidy=TRUE)
```


Load the appropriate libraries


```
if (!require("pacman")) install.packages("pacman")
pacman::p_load(plyr,readr,tidyr,dplyr,ggplot2,broom)

#library(MASS)
#library(pscl)
#library(plyr)
#library(readr)
#library(tidyr)
#library(dplyr)
#library(ggplot2)
#library(broom)
library(nlme)
```


Setup the folder with the files needed for the analysis


```
folder="~/Documents/R_Programming/LDScreen_Bioreactors"
```


Files needed:  
- genes.tab: this has the gene name (locusId), scaffold (scaffoldId), beginning position (begin), and ending position (end), systematic name (sysName), common name (name), and description (desc). tab delimited


```
genes.tab.FileName=file.path(folder,"genes.tab")

genes.tab <- read_delim(genes.tab.FileName, "\t", escape_double = FALSE, trim_ws = TRUE)
```


```
- experiment info file.
```

This has columns: Index (values are the columns in poolcount), Group (pairing structure of T0, Control, Experimental), Type (0,C, or E)


```
experimentInfo.FileName=file.path(folder,"lightdarkbioreactors_exp.csv")

expInfo<- read_delim(experimentInfo.FileName, ",", escape_double = FALSE, trim_ws = TRUE) #assumes it is comma delimited
expInfo$Group=as.factor(expInfo$Group)
```


- .poolcount file: scaffold (scaffold), position (pos). pos is followed by headers that are the experiments


```
poolCount.FileName=file.path(folder,"lightdarkbioreactors_all.poolcount.csv")

strainData<- read_delim(poolCount.FileName, ",", escape_double = FALSE, trim_ws = TRUE)
strainData$strain=1:nrow(strainData) #assign a unique Index to each strain
barcodeInfo=distinct(dplyr::select(strainData,barcode,rcbarcode,strain)) # store barcode data in its own data frame
strainData = dplyr::select(strainData,-barcode,-rcbarcode) # barcodes are removed, but strains still have their unique numbers
#Assign unique numbers to positions (same position, same scaffold) - some barcodes share the same position
temp=distinct(dplyr::select(strainData,scaffold,pos))
temp$posId=1:nrow(temp)
strainData=left_join(strainData,temp,by=c('scaffold','pos'))
rm(temp)


#Note that the Index columns don't match the experiment file.  They all have an additional SynE_ML6_set6. ahead of the Index.  Remove this
nm=names(strainData)
for (i in 1:length(nm)){
  nm[i]=gsub('SynE_ML6_set*.','',nm[i])
  nm[i]=sub("\\.","",nm[i])
}
names(strainData)=nm

startSampleNames=min(which(names(strainData) %in% expInfo$Index))
endSampleNames=max(which(names(strainData) %in% expInfo$Index))


# Determine total counts before removing strains for various criteria
# A bit complicated but it does the job
totalCounts=   gather(strainData,Index,counts,startSampleNames:endSampleNames) %>% group_by(Index) %>% summarize(total=sum(counts)) %>% mutate(ltotal=log2(total))
```


- Assign strains to genes [by position and scaffold]. Do by genes because it is shorter Note that there are many strains without a gene (insertion in intergenic regions)
- Get rid of any strains not in genes
- Get rid of strains not in the middle 80% of the gene
- Get rid of any genes without at least 3 strains


```
strainData$locusId=character(nrow(strainData))
for (g in 1:nrow(genes.tab)){
  # Which strains fall in this gene
  iStrains=which(strainData$scaffold==genes.tab$scaffoldId[g] & strainData$pos>genes.tab$begin[g] & strainData$pos<genes.tab$end[g])
  strainData$locusId[iStrains]=genes.tab$locusId[g]
}

#Get rid of strains not in genes
strainData=filter(strainData,locusId!='')

# Record the position in the gene (0 -> 1) and get rid of strains not in the middle 80%
strainData=left_join(strainData,dplyr::select(genes.tab,locusId,begin,end),by='locusId') %>% mutate(locationInGene = (pos-begin)/(end-begin)) %>% dplyr::select(-begin,-end) %>% filter(locationInGene >=.1 & locationInGene <= .9)

# Remove any genes without at least 3 posIds (not just strains)
strainData=ddply(strainData,'locusId',function(x){
  if (length(unique(x$posId))>=3){
    return(x)
  }
}) %>% tbl_df()
```


Incorporate the experimental info


```
startSampleNames=min(which(names(strainData) %in% expInfo$Index))
endSampleNames=max(which(names(strainData) %in% expInfo$Index))
strainData1=gather(strainData,Index,counts,startSampleNames:endSampleNames) %>% left_join(expInfo,by='Index')
```


Normalize within the Group by subtracting (on log scale) off the T0 data What to subtract off? Each T0 is 4x technical replicated. If we want to subtract this off, need to average these to get a single number.

Exclude genes without at least 15 T0 reads in each Group (could be spread between replicate T0s and acros strains) Then normalize by the total count to normalize for different #s of reads and then subtract off the average T0 by Group


```
T0.reads=filter(strainData1,Type=='0') %>% group_by(Group,locusId) %>% summarize(tot=sum(counts))

genes_moreThan15=T0.reads %>% group_by(locusId) %>% summarize(mn=min(tot)) %>% filter(mn>=15) %>% select(locusId) %>% .$locusId


pseudocount=1
strainData1=strainData1 %>% mutate(l2counts=log2(counts+pseudocount)) %>% left_join(totalCounts,by='Index') %>% mutate(read.normalized.l2counts=l2counts-ltotal) %>% select(-total,-ltotal)

T0.data=filter(strainData1,Type=='0') %>% select(strain,read.normalized.l2counts,Group) %>% group_by(strain,Group) %>% summarize(av=mean(read.normalized.l2counts)) 

sq.data=filter(strainData1,Type!='0') %>% left_join(T0.data,by=c('strain','Group')) %>% mutate(Group.normalized.l2counts=read.normalized.l2counts-av) %>% filter(locusId %in% genes_moreThan15) %>% select(-f,-read.normalized.l2counts,-counts,-l2counts,-av,-strand,-posId,-pos)
```


Prepare the analysis


```
# Make sure that all categorical variables are seen as such by R

# Remove any strains that don't have data in both the control and experimental. Then remove any genes that are the same
control=filter(sq.data,Type=='C')
exper=filter(sq.data,Type=='E')
commonStrains=intersect(control$strain,exper$strain)
commonGenes=intersect(control$locusId,exper$locusId)
sq.data=filter(sq.data,strain %in% commonStrains) %>% filter(locusId %in% commonGenes)

toMakeCategorical=c('scaffold','locusId','strain','Index','Group','Type')
for (columnName in toMakeCategorical){
  sq.data[,columnName][[1]]=as.factor(sq.data[,columnName][[1]])
}
```


Do the analysis

This is a linear mixed effects model. It predicts the log scaled counts as a function of a strain effect and a difference between conditions. The condition effect amounts to the difference between the averages of the experimental and control conditions. The strain effect is a random effect where strains are assumed to be a random sample of all possible strains representing a gene.

A full model (strain effect and condition effect) is fitted and compared to a model without the condition effect. The question is whether taking the condition into account actually explains the data significantly better than leaving it out. If the gene really behaves differently in two conditions, then the model that includes the condition (full) should explain the data much better than the model without it (null). A likelihood ratio test is used to test this.


```
model.full=group_by(sq.data,locusId) %>% do(fit=lme(fixed=Group.normalized.l2counts~Type,random=~1|strain,.,method='ML'))
model.null=group_by(sq.data,locusId) %>% do(fit=lme(fixed=Group.normalized.l2counts~1,random=~1|strain,.,method='ML'))

sqData.coefficient.results.full=model.full %>% tidy(fit,effects="fixed") %>% ungroup()

sqData.model.results.full = model.full %>% glance(fit) %>% ungroup() #will output a lot of red - just ignore
sqData.model.results.null = model.null %>% glance(fit) %>% ungroup() #will output a lot of red - just ignore
#sqData.obs.results = mods %>% augment(fit) %>% ungroup()

#Compare the models and do the multiple comparison
sqData.model.comparison=left_join(select(sqData.model.results.full,locusId,logLik),select(sqData.model.results.null,locusId,logLik),by='locusId') %>% mutate(neg2LL=-2*(logLik.y-logLik.x),p.value.model=pchisq(neg2LL,1,lower.tail=FALSE),p.holm.model=p.adjust(p.value.model,method='holm'),p.fdr.model=p.adjust(p.value.model,method='fdr'))


#Correct for multiple comparisons - to add other method (e.g. method x) just add  x=padj.x(p.value,method='x') in the mutate(...) block below
#sqData.Type.estimates=filter(sqData.coefficient.results,term=='TypeE') %>% mutate(padj.holm=p.adjust(p.value,method='holm'),padj.fdr=p.adjust(p.value,method='fdr'))

# Combine the model p-values and the estimates

sqData.Type.estimates.full=left_join(filter(sqData.coefficient.results.full,term=='TypeE'),select(sqData.model.comparison,locusId,p.value.model,p.holm.model,p.fdr.model),by='locusId')
```


Output the analysis


```
# output the fitted difference between experimental and control
write_csv(sqData.Type.estimates.full,file.path(folder,'LDFitness_estimates.csv'))
#write_csv(sqData.model.results,file.path(folder,'BenRubin_Set1_modelFits.csv'))
```
